# Viral diversity and dynamics, and CRISPR-Cas mediated immunity in a robust alkaliphilic Cyanobacterial consortium

**DOI:** 10.1101/2023.03.03.531066

**Authors:** Varada Khot, Marc Strous, Xiaoli Dong, Alyse K. Kiesser

## Abstract

In many industries, from food to biofuels, contamination of production systems with predators is a costly problem and requires the maintenance of sterile operating conditions. In this study, we look at the robustness of one such alkaliphilic consortium, comprised largely of a cyanobacterium *Candidatus* Phormidium alkaliphilum, to viral predation. This consortium has existed without a community crash over several years in laboratory and pilot scale environments. We look at CRISPR-Cas systems and viral dynamics in this consortium at four conditions using metagenomic analyses. Results show that while there are active viral members in this community, viral predation of the cyanobacteria is low and does not affect the community dynamics. The multiple CRISPR arrays within the Phormidium were found to be static following initial lab establishment of consortium. Multiple cryptic CRISPR-Cas systems were detected with uncertain viral protection capacity. Our results suggest that dynamics of potential viruses and CRISPR-Cas mediated immunity likely play an important role in the initial establishment of consortia and may continue to support the functional robustness of engineered microbial communities throughout biotechnology applications.

**Importance:** Biotechnology applications utilizing the function of microbial communities have become increasingly important solutions as we strive for sustainable applications. Although viral infections are known to have significant impact on microbial turnover and nutrient cycling, viral dynamics have remained largely overlooked in these engineered communities. Predatory perturbations to the functional stability of these microbial biotechnology applications must be investigated in order to design more robust applications. In this study, we closely examine virus-microbe dynamics in a model microbial community used in a biotechnology application. Our findings suggest that viral dynamics change significantly with environmental conditions and that microbial immunity may play an important role in maintaining functional stability. We present this study as a comprehensive template for other researchers interested in exploring predatory dynamics in engineered microbial communities.

## INTRODUCTION

Recent large scale viral metagenomic studies have revealed many discoveries of virus-bacteria interactions across diverse environments. For example, these studies showed the dynamics and impact of viruses on global biogeochemical cycles (1–5) and discovered new bacterial defence mechanisms (6–8), unravelling the critical roles viruses play in microbial community dynamics. Community dynamics refers to the constant microbe-microbe interplay in terms of sharing metabolites, predation and living arrangements such as biofilms. Although viruses have been recognized as important members in all microbial communities, most studies of engineered microbial systems have so far mainly focused on bacteria-bacteria (9,10) or environment-bacteria interactions (11). Engineered microbial environments differ from natural environments in that they are often closed or partially closed systems with more stable environmental conditions such as temperature, salinity, pH and nutrient concentrations, and generally lower microbial diversity. As engineered systems are designed for specific functional outcomes, loss of specific community members due to viral predation, can destabilize otherwise successful processes (12,13). Examples of this have been observed in the food fermentation industry where viral contamination has been an on-going issue (14). A recent experimental study on viruses of microalgae also showed a significant reduction in algal growth in lab cultures infected with viruses, showing potential for economic losses in this industry (15).

In the growing applications of microalgae biotechnology (16), the overall robustness of algae-based systems is of central interest. In particular, viral-associated lysis may lead to major losses of cultivated cyanobacteria (15,17). This phenomenon is commonly observed in cyanobacterial and phytoplanktonic blooms in oceans (1,18,19) and freshwater lakes (20). Yet, little is known about the extent of the impact that viruses and their dynamics have within cyanobacterial microbial communities used in biotechnology or methods to mitigate their impacts. Here we explore the dynamics of viral diversity and CRISPR-Cas mediate immunity in an alkaliphilic cyanobacterial consortium for use in biotechnology applications (21–23). A microbial consortium or community can be defined as a group of microorganisms that fulfil unique ecological roles living together. In addition to the cyanobacterium *Candidatus* Phormidium alkaliphilum, this consortium includes heterotrophs affiliated with *Bacteroidota*, Proteobacteria (Gamma and Alpha), *Verrucomicrobiota, Patescibacteria, Planctomycetota*, and Archaea (24). The consortium has displayed high productivity and robustness to environmental change, both in the lab and in larger outdoor pilot scale trials (21,25). A microbial consortium is considered robust if external perturbations do not disturb the overall function and growth of the community. The microbial community was originally sourced from alkaline Soda Lakes on the Cariboo Plateau in British Columbia, Canada (26) and has been maintained without a community collapse in a non-sterile lab environment since 2015.

Current literature on viruses from alkaline environments such as soda lakes suggests that viruses are abundant entities there and distinct to viruses from other aquatic environments. For example, tailed viruses featuring large capsids showed abundances that shifted with depth and season (27,28). Novel viruses such as those infecting alkaliphilic *Paracoccus* and giant viruses have also been recently isolated from East African Rift Valley soda lakes (29) and soda lakes in Brazil respectively (30). Additionally, high pH conditions can impact the stability of the capsid protein structure (31,32), and may pose a unique evolutionary pressure for viruses in high pH, high salt environments.

Studying viral dynamics within highly diverse microbial communities poses many challenges because of the added complexity of strain-level diversity and the multitude of microbial interactions that are present. Challenges associated with viral sequence identification, historically low rates of host predictions to the genus level (33) and the magnitude of the network connecting multiple viruses associated with multiple hosts (34) make spatiotemporal dynamics difficult to follow. As such, less diverse microbial consortia that exist as established communities or consortia make a more tractable system to study viral community dynamics as microbial populations and dynamics can be tracked over time and under a variety of experimental conditions. In the context of biotechnology, such a model system could be useful to effectively predict functional outcomes and community robustness.

To characterize alkaliphilic viral diversity in this engineered system, we used metagenomes of the consortium previously obtained at high (pH 10-11) and low (pH 8-9) pH and with three different nitrogen sources (ammonia, nitrate, and urea) (23,24). A consensus approach was used to identify DNA sequences associated with viruses (35), classify their taxonomy (36) and to infer their hosts (37) from these metagenomes. To explore the bacterial defenses against viruses, we focused our analyses on CRISPR-Cas (Clustered Regularly Interspaced Short Palindromic Repeat) systems. CRISPR-Cas systems are a microbial adaptive “immune system” present in 36% of bacteria and 75% of archaea (38). They carry a history of interactions with viruses and mobile genetic elements in the form of ‘spacers’ – snippets of 25-35 base pairs inserted into the CRISPR array, which can be useful for host predictions. CRISPR arrays are typically located adjacent to CRISPR-Associated (Cas) proteins, which are required for the systems to be functional and therefore also important to identify.

## METHODS

### Dataset

Eleven metagenomic datasets were used for this study as described in Supplementary Table 1. These included four metagenomes from the soda lakes (GLM, DLM, PLM, LCM) (26), six metagenomes of the laboratory photobioreactor cultures (Year 0, Year 1, Year 2) (23,24) and one complete genome of *Candidatus* Phormidium alkaliphilum (23). In all analyses, the Year 1 metagenomic assembly was used as a reference point to compare other datasets to. Metagenomic reads from the consortium grown under four experimental conditions in a photobioreactor described by Ataeian et. al. (23,24) (Supplementary Table 1), were used to create the Year 1 assembly. These conditions, varied by pH and nitrogen source, consisted of low-pH ammonium, low-pH nitrate, high-pH nitrate, and high-pH urea. The total number of paired-end reads from each metagenome was between 40-45 million and the average read length was 145-150 base pairs. Reads from the four conditions were co-assembled using MEGAHIT (34) and are referred to “Year 1 metagenome”. The resulting contigs were binned into Metagenome-Assembled-Genomes (MAGs) using MetaBAT v2.12.1 (35). These MAGs are the same as referenced in Ataeian et. al. (24).

### CRISPR Array and *Cas* gene Identification

#### CRISPR Array Identification

CRISPR arrays were identified from Year 1 metagenomic reads and assembled contigs using Crass (39) and MinCED (40) respectively, both with default settings. Reads identified by Crass as belonging to CRISPR arrays were mapped to contigs using BBMap (41). An in-house script was then used to retrieve the taxonomic associations for the MAGs that contained the CRISPR reads. Crass identified CRISPR arrays were compared to those identified by MinCED by the associated MAG and number of spacers in each array using an in-house script. These arrays were then dereplicated to keep only 1 unique set of CRISPR arrays per MAG.

#### CRISPR Array Comparison

CRISPR arrays from the Year 1 co-assembly were then compared to other consortium metagenomic assemblies (Year 0, Year 1 and Year 2) and assembled metagenomes of microbial mats from soda lakes on the Cariboo Plateau (GEM, PLM, DLM, LCM) (Supplementary Table 1). using Blastn (42) with the options “ -task blastn-short -outfmt 6”. CRISPR arrays from bin00 (*Candidatus* Phormidium alkaliphilum MAG) from the Year 1 assembly was also compared to the *Candidatus* Phormidium alkaliphilum whole genome sequence (Year 3) using Blastn (same parameters as above). Blast results were then filtered for 100% identity and 100% query cover to find exact matches of spacers over time within the consortium.

#### Cas Gene Identification

Cas genes were annotated in the Year 1 co-assembly using the annotation pipeline MetaErg (43), which uses Hidden Markov Models (HMMs) to identify Cas genes (44). Identified clusters of Cas genes were then classified as specific “Cas systems” according to Makarova et al. (45).

#### Viral Sequence Identification

Potential viral DNA sequences within the contigs from the Year 1 assembly were identified using Virsorter2 (35) with the options “–keep-original-seq –include-groups dsDNAphage,ssDNA –min-length 5000 –min-score 0.5 all”. The resulting 213 viral contigs (VC) were then quality-checked, and host genes trimmed from the ends of identified contigs using CheckV (46). The VCs were filtered using criteria in Viral Sequence Identification Standard Operating Procedure (SOP (47)) and only contigs in the “keep” category with a sequencing depth greater than or equal to 5 in at least 1 sample were kept for downstream analyses. The final number of viral contigs was 83. Dramv (48) was used to annotate genes on these 83 contigs. Reads from all other metagenomic datasets were mapped to the viral contigs identified in the Year 1 assembly using BBMap (41) to trace the origin and temporal dynamics of potential viruses.

#### Viral Phylogenomics and taxonomy

The viral contig dataset was then compared to reference viral genomes from NCBI Viral Refseq Release 208 (2021) (49) using Blastp (42). The bitscores of these matches were used to calculate a Dice coefficient (50,51) to measure the distances between the viral contigs and reference viral genomes. Based on Dice coefficients, sequences were clustered using neighbour-joining to generate a phylogram showing relationships between viral contigs from the consortium and reference viruses. In parallel, contigs were also taxonomically classified using VConTACT2 (36), which uses a reference database of archaeal and bacterial viruses from Viral Refseq v86. The top classification for each viral contig from VConTACT2 was retained.

#### Virus-Host Predictions

Four methods were used to determine the most probable host for a given viral contig. 1) Previously identified CRISPR spacers from binned metagenomic contigs were dereplicated and aligned to the viral contigs using BLASTn with the options “-task blastn-short -dust no -outfmt 6” (42). Results were filtered for <= 2 mismatches, alignment length > =20bp and an evalue <= 10^-5^. 2) Viral contigs were aligned to the entire metagenomic assembly using BLASTn and alignments > 100bp were considered for host prediction. 3) Tetranucleotide frequency profiles between viral contigs and MAGs were calculated using an in-house script and the cosines of the angle between the tetranucleotide vectors were used to determine proximity of vectors. Top two matches per VC, scoring >= 0.95 were retained for host prediction. 4) read depth profiles of viral contigs and host genomes were generated using bbmap to map reads from each sample to the CC2018 assembly and providing the subsequent .bam files to CheckM coverage (default settings) (52). These sequencing depth profiles were then clustered using hierarchical clustering with a “Bray-Curtis” distance matrix and an “average” clustering algorithm in Python, resulting in a dendrogram of the viral contigs and MAGs. An in-house script was used to parse the dendrogram to find the closest match between each VC and MAG. Viral contigs clustering closely with MAGs were identified as potential virus-host pairs. All four methods for host predictions were used in conjunction to identify a “most probable host” for each viral contig.

## RESULTS

### 1. HOST ADAPTIVE IMMUNITY IN THE CONSORTIUM

CRISPR arrays detected by Crass (39) and MinCED (40) were found in 18 out of 29 metagenome-assembled genomes (MAGs) from the Year 1 consortium metagenome, sampled after one year of laboratory enrichment (Figure 1). Spacers from the Year 1 CRISPR arrays were compared to previous metagenomes using BLASTn (42) from the enrichment time points (Year 0, Year 2, Year 3) and alkaline soda lakes (GEM, PLM, DLM and LCM, Supplementary Table 1). The number of spacers within the genome of *Cand*. P. alkaliphilum appears highly consistent across multiple years within the consortia, suggesting these spacers have remained static throughout multiple years of enrichment. When compared to the lakes, spacers from *Cand* P. alkaliphilum CRISPR arrays were only found in the GEM and PLM samples, although the original consortium was seeded from all four soda lakes. The increase in spacers from the original lakes to the enrichment indicates that the cyanobacteria acquired new spacers in all CRISPR arrays (except CRISPR5) during the initial adaptation to the laboratory environment and these have been carried forwards over multiple years.

**Figure 1:**
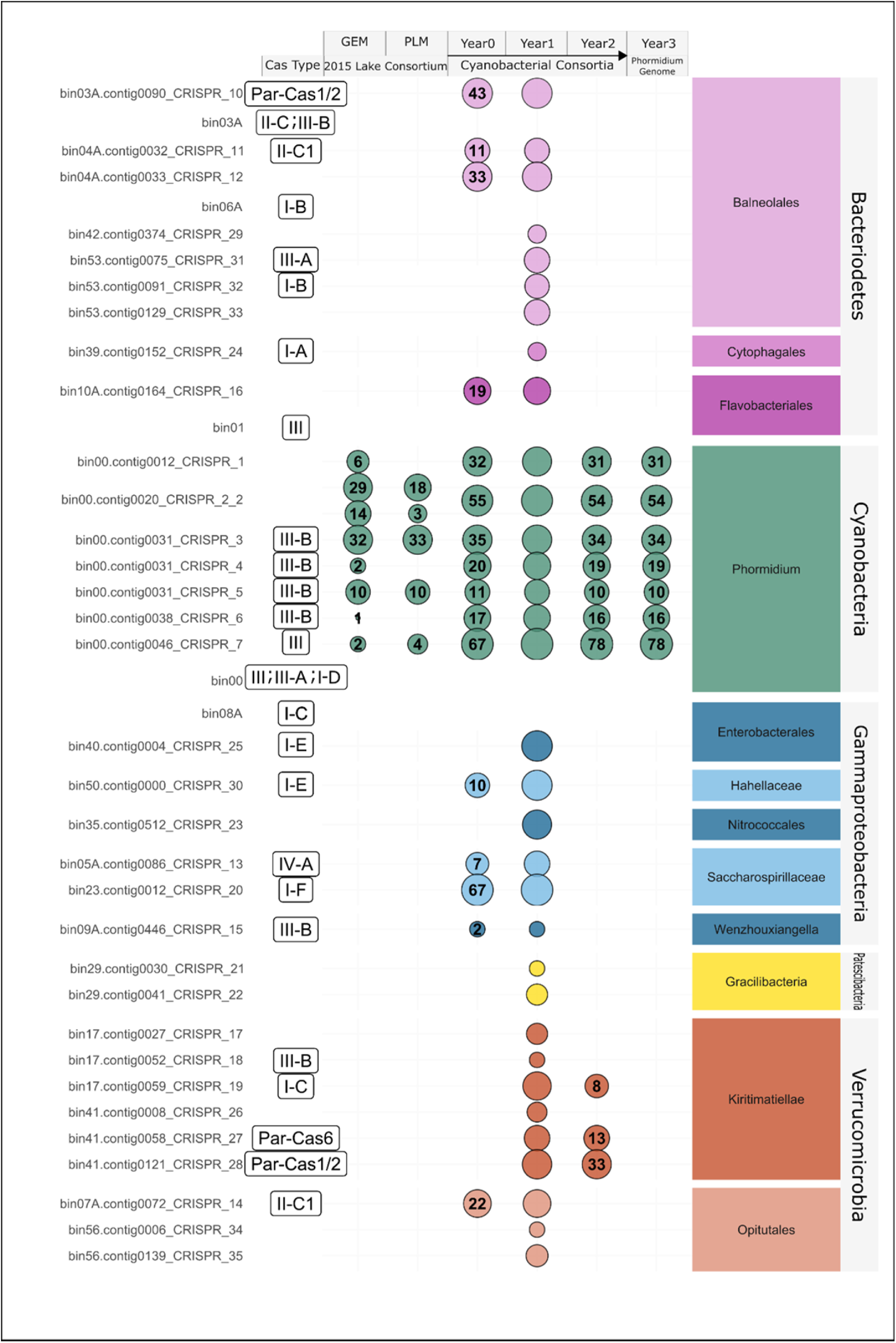
CRISPR arrays, Cas-system types and spacers from the Year 1 reference metagenome MAGs. Each line shows a single CRISPR array and associated Cas proteins. For each MAG and metagenome, the number within the bubble indicates the number of CRISPR spacers matched to the Year 1 reference metagenome.

To further explore the potential origin of these spacers, spacers from *Cand* P. alkaliphilum CRISPR arrays were also searched against the Year 1 assembly using BLASTn (“-task blastn-short”). Conclusive matches from this BLAST search (100% identity and (e-value < 1E-3) were only to unbinned contigs, for which taxonomy could not be determined and did not include any viral proteins or contigs. The same spacers were also searched against *Cand* P. alkaliphilum whole genome (Year 3) to look for self-targeting spacers using BLASTn (short task). No conclusive matches for self-targeting spacers were found as they were either short in length (length < 19bp) or with high e-values (e-value > 1E-3).

For the heterotrophic consortium members, CRISPR arrays were more dynamic during three years of laboratory cultivation and CRISPR arrays were not conserved between time points of laboratory cultivation. This could be most easily explained by rapid turnover of organisms in the community, rather than a turnover of CRISPR spacers, as MAGs from Year 1 were not found in Year 0 or Year 2 metagenomes. High turnover of heterotrophic consortium members was also reported previously for this consortium (24).

As CRISPR-Cas systems often function along side associated *Cas* genes, Cas genes were identified in the MAGs from Year 1 using Hidden Markov Models (44). Cas genes were found adjacent to 20 out of 33 CRISPR arrays (Figure 1). Thirteen out of 31 MAGs had no CRISPR arrays or *Cas* genes.

Type I and Type III Cas systems were the most common in the consortium. These are also the most commonly found Cas systems in bacteria, with genes for both systems often found in the same genome (53,54). This was also the case in our dataset where *Cand*. P. alkaliphilum and a species of Kiritimatiellae (Figure 1, bin17), each contained both Type I and Type III systems. Overall, the presence of CRISPR-Cas systems in most of the consortium members would be consistent with the robust nature and long-term stability of our system overall. Yet, the observed high turnover in heterotrophic community members (24) suggests multiple factors that may be at play in the modulation of the heterotrophic populations.

### 2. VIRAL DIVERSITY AND DYNAMICS IN THE CONSORTIUM

A survey of the Year 1 metagenome for potential viruses with VirSorter2 (35) characterized the viral abundance and diversity in the consortium. The identified contigs were filtered based on their length (>5 kb) and sequencing depth (>5x in any sample), resulting in 83 identified viral contigs (VCs). Twenty-six of the 83 VCs could be taxonomically classified using VConTACT2 (36), with 25 out of 83 viral contigs affiliated with the order Caudovirales (Supplementary Table 3). Sequencing depth of the viral contigs was used to differentiate viral populations in each of the samples (Low-pH Ammonia, Low-pH Nitrate, High-pH Nitrate, High-pH Urea) contributing to the Year 1 co-assembly. A mean sequencing depth of the VCs was within the same order of magnitude of the heterotrophic MAGs (Supplemental Table 3) at 15.32, 14.90, 7.61 and 2.86 in the Low-pH Ammonia, Low-pH Nitrate, High-pH Nitrate, High-pH Urea samples respectively. The notable exception was uVC86 with a sequencing depth an over order of magnitude higher than the mean (Figure 2).

**Figure 2.**
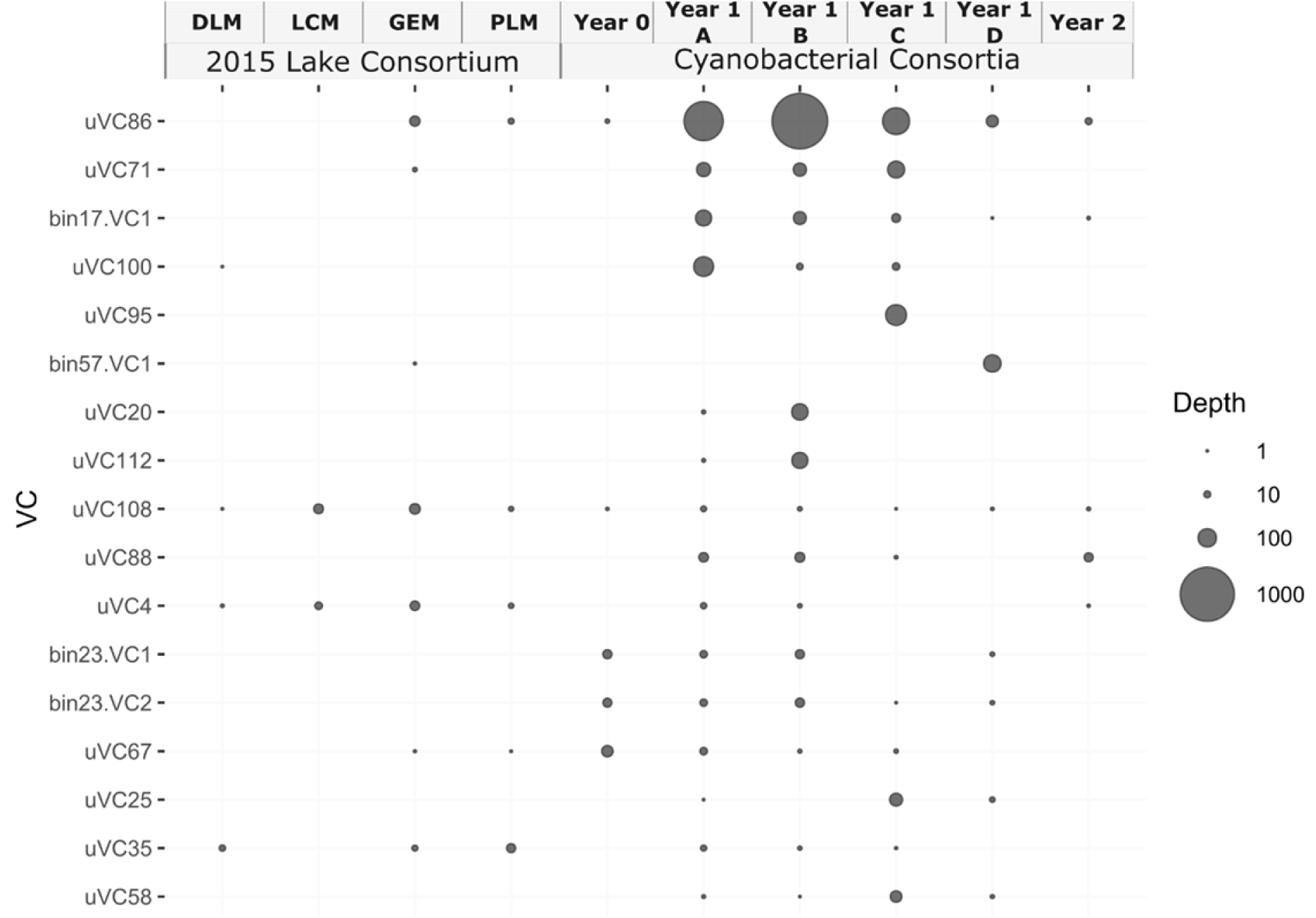
Viral Contig abundance. Numbers of reads mapped to the seventeen most abundant viral contigs (VCs) from all studied metagenomes. For year 1, viral contigs were found in the Year 1 metagenome from the following samples: A) Low-pH Ammonia, B) Low-pH Nitrate, C) High-pH Nitrate, D) High-pH Urea. Abundances of all 83 VCs are presented in Supplementary Figure 2.

To explore the potential relationship of the viral particles with the consortium community, hosts were computationally assigned to the VCs by four methods: 1) matching CRISPR spacers, 2) high sequence similarity 3) similar tetranucleotide frequencies and 4) similar read depth profiles across samples. Sixty VCs had host predictions using at least one method (Figure 3, Supplemental Table 4) (55). MAGs identified as potential hosts included a range of heterotrophic bacteria from the community, affiliated with *Alphaproteobacteria, Bacteriodia, Gammaproteobacteria, Phycisphaerae, Rhodothermia* and *Verrucomicrobiae* (Figure 3). Although many host MAGs possessed CRISPR arrays, no CRISPR spacers matched to their associated VCs at high confidence (100% identity and e-value <= 10^5^). This indicated that the information stored in bacterial CRISPR arrays was likely left over from older, no-longer-relevant viral infections or had a non-viral origin (56). Overall, the host prediction methods used here in parallel linked VCs to multiple hosts (Figure 3), rather than linking each VC with a specific organism in the community. This interpretation of host prediction, while broad, reflects the potential of current computational methods within this dataset. Therefore, in cases where multiple prediction methods agreed on the host, it resulted in a more robust overall virus-host linkage.

**Figure 3.**
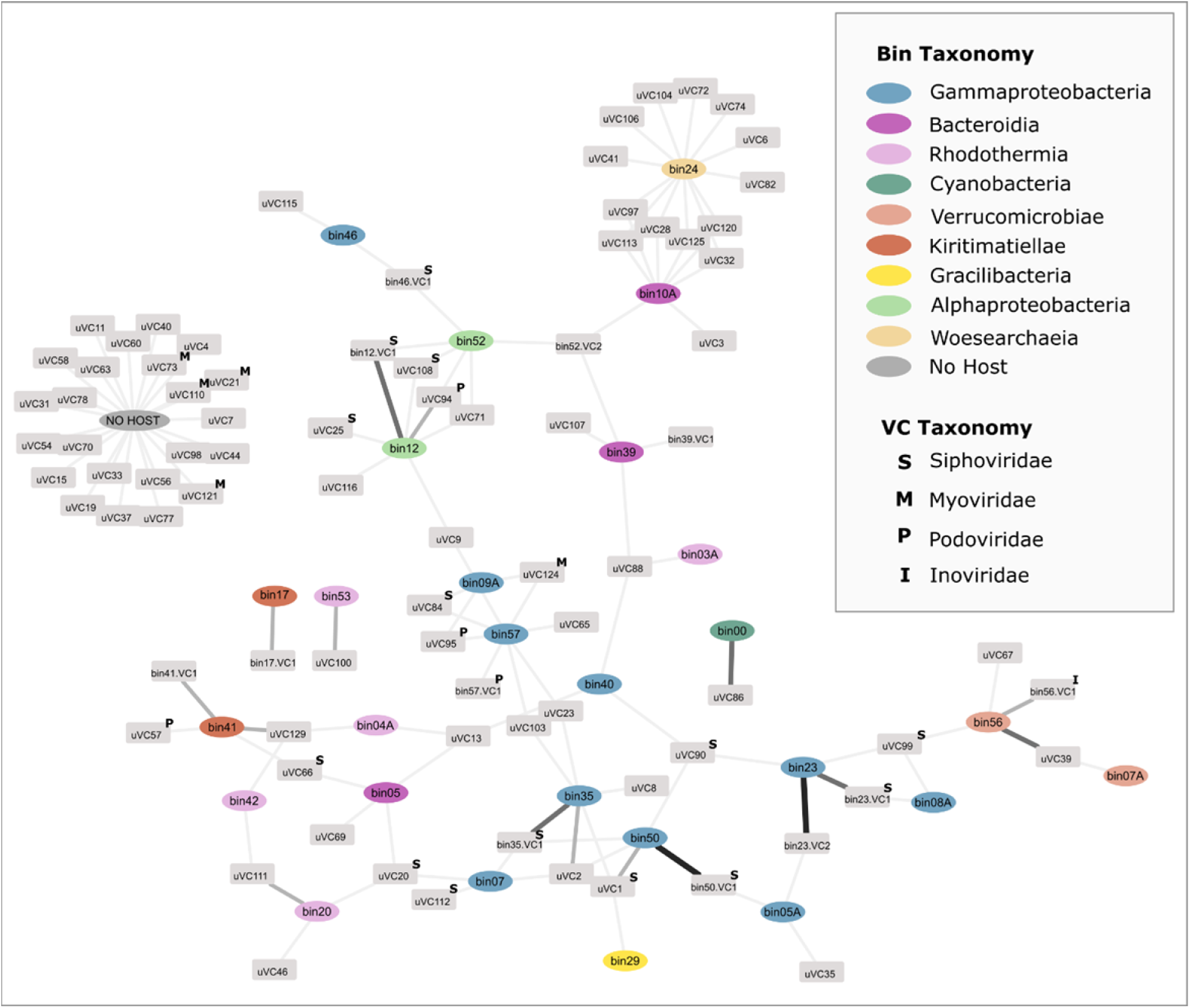
Network of viral contigs (VCs) linked to computationally predicted hosts. The width and colour of line represent the confidence of the host association. VCs with no associated hosts are clustered together. MAGs are coloured based on taxonomy at the class level.

For sixteen VCs, a robust virus-host linkage was determined with two or more methods agreeing on the host prediction. This included 7 VCs that were binned with the host contigs. Binning criteria of MetaBAT2 (35), used for binning, included tetranucleotide frequencies and sequencing depth profiles to group contigs into a bin. Thus, it was unsurprising that binning worked as a host prediction method, regardless of whether the virus was integrated into the host genome or not. In four cases (bin12.VC1, bin23.VC1, bin23.VC2 and bin41.VC1), host genes were found flanking the VC, indicating that it was, indeed, a prophage (Supplemental Table 4, Tab 2). Conversely, for VCs bin35.VC1, bin50.VC1 and bin75.VC1, it was not possible to determine whether these were prophages or free viral fragments as no host genes were found on the same contig. Sequencing depth profiles of bacterial MAGs and viral contigs were also compared to investigate whether viruses and their hosts follow a similar pattern of abundance in the different experimental conditions (Supplemental Figure 2). This method provided host predictions for 27 viral contigs. Generally, the sequencing depth between bacterial contigs and viral contigs within the same MAG co-varied across conditions, indicating that the VC might have originated from a prophage (Supplementary Figure 2), though the viral contigs had lower overall sequencing depth than their identified potential hosts. This could be interpreted that the VC originated from a virus (temperate or lytic) that only infected a portion of the host population. Future efforts to further explore the temporal dynamics of these virus host interactions could include a time series study with closely spaced time intervals or perturbations to conditions and could yield significant insights to how theses interactions integrate into the overall robustness of the system.

Several studies have documented the existence of CRISPR-Cas systems and integrated phages in a single genome (54,55,57). One such study found that bacteria are likely to have self-targeting spacers if they also have an integrated phage, although this was not found to be the case in our study.

One viral contig of particular interest was bin57.VC1, which was binned with contigs from its host population a Gammaproteobacteria classified as a species of Wenzhouxiangella. This VC only appeared in the high pH-urea condition with 11x higher sequencing depth than the identified host of bin57. The viral contig Bin57.VC1 was relatively large (∼41kbp) and did not contain bacterial genes on either end. This indicated that bin57.VC1 likely originated from an active virus in the lytic cycle in the high-pH urea conditions. Although binned, bin57.VC1 did not have hits to any other Wenzhouxiangella genomes or viruses, when searched using Blastn. DRAMv annotations identified genes for both putative capsid and head-tail connecter proteins (Supplemental table 2). Based on previously studied populations of Wenzhouxiangella from high-pH environments, this organism is considered to have a predatory lifestyle, with genes for proteolytic activities and lantibiotic biosynthesis pathways (21,55). Previous stable isotope probing and proteomics of the consortium suggest the bin57 Wenzhouxiangella uptakes macromoleucles from *Cand*. P. alkaliphilum. An activate viral infection of this predatory organism may result in an increase in dissemination of fixed carbon from the Cyanobacteria to the heterotrophic community members (Supplementary figure 2). Indeed, viral activity is known to play a significant role in carbon dynamics on a global scale (18). Additional studies of the dynamics of Bin57.VC1 within the consortia could explore the role of the virus and carbon cycling and its potential to disrupt the overall stability of the consortia in biotechnological applications.

### 3. ROBUSTNESS OF *CAND*. P ALKALIPHILUM TO VIRAL PREDATION

As the cyanobacterial consortium depended on the primary productivity of *Cand.* P. alkaliphilum (22,23), its robustness against viral predators was further investigated. We found significant evidence for adaptive immunity in the genome in the form of seven CRISPR arrays and five clusters of Cas genes (Figure 1). As mentioned above, it appeared that spacers in some of these arrays were acquired during initial adaption to the lab. After the initial adaptation, the lack of change in CRISPR arrays of the *Cand.* P. alkaliphilum (Figure 1) could be attributed to the lack of new viral predators in the laboratory environment, or the presence of other, more effective defence mechanisms that prevented activation of the CRISPR-Cas systems and subsequent incorporation of spacers. Alternatively, the cyanobacterium’s CRISPR systems may serve a different, unknown purpose unrelated to viral predation and that does not require updating the arrays (58). For example, given partial spacer self-targets, CRISPR-Cas systems may be involved in gene regulation, DNA repair or programmed cell death (58,59).

Clusters of Cas genes found in the *Cand.* P. alkaliphilum genome belonged to type I and III systems and included a complete type III-B Cas system with a reverse transcriptase adjacent to a Cas1 gene and three CRISPR arrays. Other Cas systems in this genome included a type I-D, with no associated CRISPR array and missing adaption modules (Cas1 and Cas2 genes) as well as a type III-B with a CRISPR array but a missing expression module (Cas6). The *Cand.* P. alkaliphilum genome also contained a ‘hybrid’ type III system with Cas genes from A, B and D subtypes, including all the modules needed for a functional system (CRISPR array plus adaption, expression, and interference modules (45)). Other Cas genes, Cas3 and Csa3, were also present, with no association with the CRISPR-Cas loci. Multiple and incomplete Cas systems with missing Cas1 and Cas2 genes have been discovered in previous surveys of Cyanobacteria (60).

Type I and type III systems, both found in the Phormidium genome, have different modes of defense via spacer acquisition and target DNA cleavage (61–63) and may be providing synergistic immunity through redundant systems (54) or by sharing Cas modules and CRISPR arrays. Type III Cas systems have broad target specificity as they do not use protospacer adjacent motifs (PAMs) to recognize cleavage sites like type I and II systems, leading to the cleavage of even partial target matches (63). A combination of type I (specific) and type III (broad) Cas systems may ensure that viruses that escape the type I system are captured by the type III system, particularly if CRISPR arrays are shared between the systems. This in part may explain the lack of the spacers with viral sequence targets found in consortium metagenomes as these viruses may have become extinct during the transition from the lakes. In the case of the Phormidium, the type I-D system may be using CRISPR arrays or adaption modules *in trans* from the type III systems as no CRISPR arrays were closely associated with the I-D loci (45). Type I and type III systems are often observed together in genomes and are speculated to share Cas genes (particularly Cas1, Cas2, Cas6) (53,56,60,64). The diverse Cas systems also ensure adaptive immunity to double stranded DNA (all systems), single stranded RNA (type III-B), and single stranded DNA (type I-D effector Csc2). Orphan CRISPR arrays, present in the Phormidium genome, are arrays which are not adjacent to Cas genes and are often found in genomes with multiple CRISPR-Cas systems. The functional significance of these arrays is not well understood; however, some Cas systems are able to use orphan CRISPR arrays in *trans*. Orphaned CRISPR arrays could be a result of the loss of Cas genes, the formation of a new CRISPR array or an array inserted by transposable elements (65). Several recent studies show that multiple remote CRISPR arrays are still functional and could play a role in long-term infection memory (66,67). CRISPR dormancy could be beneficial for evolving the genome through DNA uptake and competency (59). Further studies which specifically test the functionality of the CRISPR arrays to determine whether the unchanging CRISPR arrays in *Cand*. P. alkaliphilum are dormant, have been orphaned or perform other cellular functions. These could include, removal of the CRISPR arrays or Cas genes, insertion of Cas genes into another organism or applying viral predatory pressure.

In addition to adaptive immunity, we also looked specifically for viral contigs which came from viruses that infected *Cand*. P. alkaliphilum MAG (bin00). Based on the “Kill-the-Winner” theory (68), we expected the Phormidium to have many viral predators as it was the most abundant organism in the community at 80-90% abundance. However only one viral contig (uVC86) matched with the *Cand.* P. alkaliphilum during host prediction (Figure 2). Other viral predators may have been RNA viruses, which were likely missed during DNA extraction. The virus-host association between the Phormidium and uVC86 was supported by sequencing depth profiles (Figure 4A), tetranucleotide frequencies and the presence of identical sequence fragments (Figure 4C).

**Figure 4.**
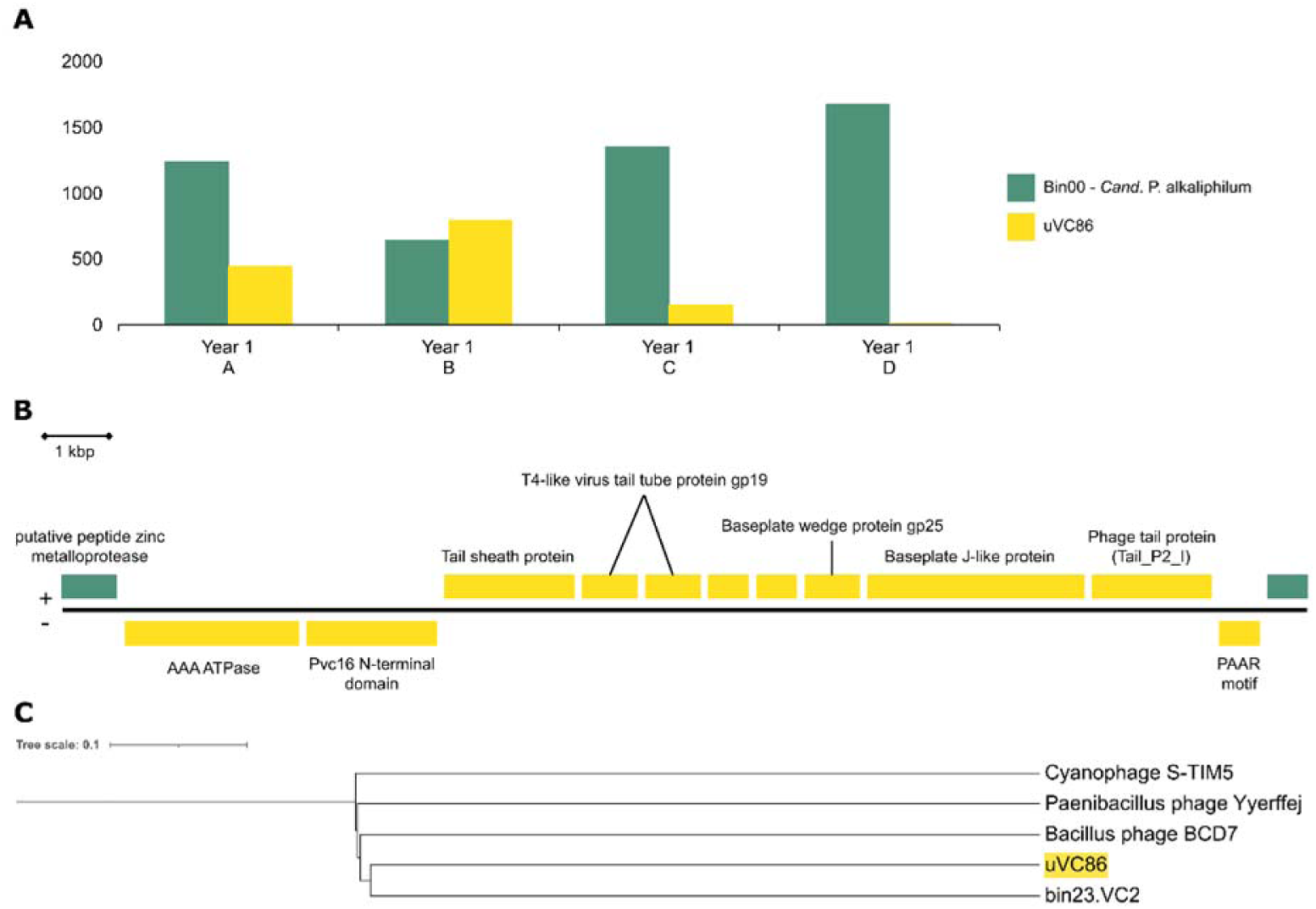
A) Sequencing depths of *Cand*. P. alkaliphilum and uVC86 in four experimental conditions in Year 1 of growth for Y1 A (Low pH-Ammonia), Y1 B (lLow pH-Nitrate), Y1 C (High pH-Nitrate), Y1 D (High pH-Urea. B) Placement of uVC86 with its closest relatives from RefSeq v208 (49) and the VCs from this study on a neighbor-joining tree calculated from Dice distances. C) Genes on contig uVC86 annotated by Dramv (48) in yellow and cyanobacterial genes in green.

However, despite the large number of Phormidium CRISPR spacers present, no spacers matched uVC86. This relatively small viral contig, of only 13kbp, had the highest abundance of all the viral contigs. The metagenomes from Good Enough and Probe lakes contained a low number of reads that mapped to uVC86 (26 and 7respectively) (Figure 2). Reads mapping from the lakes covered ∼70% of the uVC86 contig from the enrichment, suggesting the lakes as the origin for this viral sequence. This indicated that any potential viruses representing uVC86 found in the consortium at least partially originated in the lakes. The Year 1 metagenomes contained hundreds of reads that mapped to uVC86 and interestingly, abundances of uVC86 and *Cand*. P. Alkaliphilum in the four experiments were negatively correlated (Figure 4A). While the cyanobacterium’s abundance was highest in the two high pH experiments, uVC86’s abundance was highest in the two low pH experiments and very low or below detection limit at high pH. Anticorrelation of abundances of the viral contig and its putative host might, for example, be explained by 1) Successful viral infections in the low-pH conditions leading to increased uVC86 and decreased *Cand*. P. Alkaliphilum abundance; 2) Destabilization of viral (capsid) proteins in the two high pH experiments, leading to lower uVC86 abundance and success (32).

To identify potential relatives of uVC86, we compared the encoded proteins to those encoded on the other viral contigs as well as all Viral Refseq proteins, via Blastp. The resulting Dice-distance based phylogenetic reconstruction showed the closest similar protein sequences found for uVC86 was bin23.VC2, a viral contig predicted to have a different host within the same consortium, a Saccharospirillum species (Figure 4B). The Dice-distance phylogenetic reconstruction was based on homology of individual predicted protein coding sequences, allowing for rearrangement of protein coding sequences rather than overall nucleotide identity which requires gene order to be conserved (69). Despite being neighbours in this tree, viral contigs uVC86 and bin23.VC2 did not appear to be closely related, having an average amino acid identity of only 12% between shared proteins, when compared using Blastp (42). Taxonomic assignment for uVC86 with vConTACT2 was also not possible. Thus, it appeared that uVC86 is a unique cyanophage with no closely related known viruses.

Functional gene annotations by DRAMv (48) provided some clues about this potential virus’s lifestyle and taxonomy (Figure 4C, Supplementary Table 3, Tab 2). Several genes were shared with T4-like phages (tail tube and baseplate wedge proteins), and some had distant similarity to P2-like phage proteins (baseplate J-like and phage tail). Both these viruses are affiliated with Myoviridae. uVC86 encoded no capsid, integrase, or replication genes, so it is likely an incomplete viral genome, unable to replicate by itself. Viral contigs of comparable sequencing depth that could potentially complement these missing genes in uVC86 were not found. Furthermore, both ends of uVC86, encoded putative host genes (bin00.contig 4394, gene 1 and gene 12, Supplementary Table 3, Tab 2) flanking the cluster of viral genes (Figure 3C). One of these genes was a M50 metallopeptidase MER0890051; MEROPS peptidase database (70), while the other was of unknown function. Both genes had high similarity to genes of a Geitlerinema species, a close relative of *Cand*. P. alkaliphilum. As there was no evidence of Geitlerinema in the consortium, we assumed that these genes originated from *Cand*. P. alkaliphilum, even though, they were not present in the (circular) whole genome sequence of *Cand*. P. alkaliphilum (23). They most likely originated from a *Cand*. P. alkaliphilum subpopulation. Thus, it appears that uVC86 was a remnant of a prophage present in the genome of a subpopulation of Cand. P. alkaliphilum, a subpopulation that was enriched at low pH. The differences in uVC86 abundances between experiments were likely caused by differences in abundances of the host subpopulation. Coexistence of multiple Cand. P. alkaliphilum subpopulations with different ecological fitness, dependent on pH and nitrogen source, was previously reported and is a key aspect of the robustness of the consortium. The existence of a remnant of a potential phage genome within these subpopulations is consistent with a dynamic nature of phage-host dynamics within natural environments.

## Conclusion

From a biotechnology perspective, success of an autotrophic-based consortium is reliant on the survival of the primary producer, in this case *Cand*. P. alkaliphilum. Previous research has demonstrated the consortium’s robustness in the context of environmental change (21,24), as well as the different ecological roles filled by the heterotrophic members of the consortia and the characteristics of *Cand*. P. alkaliphilum. Here, we explored further the potential robustness of the consortium to viral predation. Many consortium members, including the *Cand*. P alkaliphilum, displayed adaptive immunity in the form of one or more CRISPR-Cas systems. Based on the diversity and abundance of its Cas genes, we expect the Phormidium to have comprehensive adaptive immunity that targets both RNA and DNA viruses. However, the observed lack of dynamicity of the CRISPR arrays points to greater complexity of purpose for these systems. The survey of potential viral predators in the consortium highlighted the presence of several diverse putative viruses that exist in this high pH system. The recovery of only one partial viral genome of a potential cyanophage, despite the high abundance of *Cand*. P alkaliphilum, suggests an insufficient predatory pressure to maintain cyanobacterial innate immunity within the established consortia. The addition of new CRISPR spacers within the first year of the consortia being established strongly suggests a critical role of adaptive immunity of *Cand*. P. alkaliphilum populations in the *initial establishment* of the robust consortium.

## Acknowledgments

We would like to thank Maryam Ataeian and Jackie Zorz for providing access to and help with sequencing data.

